# Forcing contacts between mitochondria and the endoplasmic reticulum extends lifespan in a *Drosophila* model of Alzheimer’s disease

**DOI:** 10.1101/706408

**Authors:** Juan Garrido-Maraver, Samantha H. Y. Loh, L. Miguel Martins

**Affiliations:** MRC Toxicology Unit, University of Cambridge, Lancaster Road, Leicester LE1 9HN, UK

**Keywords:** *Drosophila*, organelle contacts, mitochondria, endoplasmic reticulum, Alzheimer’s disease

## Abstract

Eukaryotic cells are complex systems containing internal compartments with specialised functions. Among these compartments, the endoplasmic reticulum (ER) plays a major role in processing proteins for modification and delivery to other organelles, whereas mitochondria generate energy in the form of ATP. Mitochondria and the ER form physical interactions, defined as mitochondria-ER contact sites (MERCS) to exchange metabolites such as calcium ions (Ca^2+^) and lipids. Sites of contact between mitochondria and the ER can regulate biological processes such as ATP generation and mitochondrial division. The interactions between mitochondria and the ER are dynamic and respond to the metabolic state of cells. Changes in MERCS have been linked to metabolic pathologies such as diabetes, neurodegenerative diseases and sleep disruption.

Here we explored the consequences of increasing contacts between mitochondria and the ER in flies using a synthetic linker. We showed that enhancing MERCS increases locomotion and extends lifespan. We also showed that, in a *Drosophila* model of Alzheimer’s disease linked to toxic amyloid beta (Aβ), linker expression can suppress motor impairment and extend lifespan. We conclude that strategies for increasing contacts between mitochondria and the ER may ameliorate symptoms of diseases associated with mitochondria dysfunction.

## INTRODUCTION

Eukaryotes include several unicellular and multicellular organisms including fungi, plants and animals. Eukaryotes differ from prokaryotes in that they have a compartmentalised cytoplasm with membrane-bound organelles. Organelles give eukaryotic cells several advantages, including providing special environments that favour specific biochemical reactions required for life. In eukaryotic cells, the endoplasmic reticulum (ER) provides an environment for protein and phospholipid synthesis and the storage of ions such as Ca^2+^. Mitochondria are responsible for the conversion of energy stored in the chemical bonds of nutrients into ATP, the cellular energy currency. Organelles have been studied in isolation, but now it is clear that they also form contacts. The most well-characterised organelle contact sites are formed between the ER and mitochondria (reviewed in ^1^). Sites of contact between mitochondria and the ER regulate lipid synthesis, Ca^2+^ signalling, mitochondrial energy generation and mitochondrial division.

Alterations in contacts between mitochondria and the ER can cause mitochondrial calcium overload and an increase in mitochondrial reactive oxygen species (ROS), and such alterations in the liver have been linked to obesity, a medical condition, in mice^2^. In neurons, alterations in contacts between mitochondria and the ER can result in neuronal cell death^3^, defects in the release of synaptic vesicles^4^ and ER stress^3, 5^.

The connections between mitochondria and the ER have been manipulated in cultured cells and increasing such links can commit cells to a death pathway^6^. Synthetic linkers that increase contacts between the ER and mitochondria in the mouse liver induce a transient induction of ER stress that resolves over time ^2^.

Here, we report a *Drosophila* model for expressing a synthetic linker between the mitochondria and the ER. We used this *in vivo* model to explore the long-term consequences of forcing contacts between these two organelles. We showed that flies in which the contacts between mitochondria and the ER are enhanced have a fragmented mitochondrial network and increased levels of ROS. The sustained expression of an artificial linker also leads to enhanced motor activity and extended lifespan. We also showed that the improved health span conferred by the expression of this artificial linker counteracts the toxic effects that result from the expression of a toxic version of amyloid-β (Aβ) that causes Alzheimer’s disease in humans.

## RESULTS

### Tethering ER and mitochondria in *Drosophila* using an engineered linker

The association between mitochondria and the ER has been widely documented in different experimental models. The study of this association has been the focus of intensive research, using both cellular and animal models. However, the diversity of the reported functions of these contacts suggests that their role is still unclear (reviewed in ^7^).

Here, we used a genetic approach to explore the effects of tethering the ER and mitochondria in fruit flies (*Drosophila melanogaster*) using an artificial linker. To increase the physical coupling between both organelles, we used a previously reported linker that induces enhanced proximity (6 nm) between mitochondria and the ER in mammals ^6^. We first engineered a codon-optimised version of this synthetic linker for expression in *Drosophila*. This synthetic linker (from here on referred to as “linker”) consists of monomeric red fluorescent protein (RFP) with a mitochondrial targeting sequence at the N terminus and an ER targeting sequence at the C terminus (Fig. 1a). Transgenic flies expressing this construct were generated by PhiC31 integrase-mediated transgenesis (see methods) and the expression of the linker was assayed by immunofluorescence. Confocal analysis of the larval wing discs revealed punctate staining of the linker (Fig. 1b).

**Figure 1.**
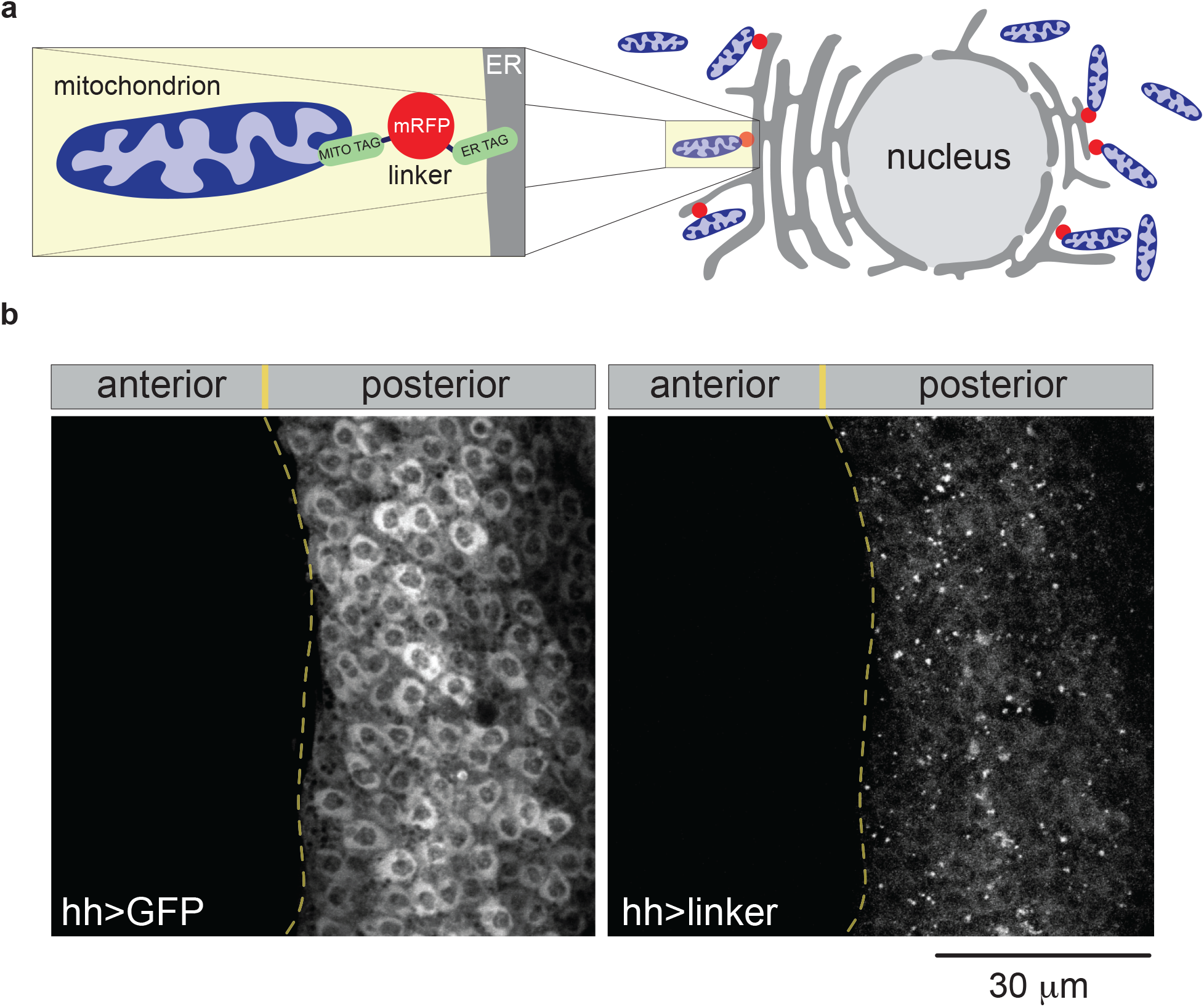
*In vivo* expression analysis of an engineered tether between mitochondria and the ER. (**a**) A schematic of the linker tethering mitochondria (blue) and the ER (grey). The magnified area on the left details the relative position of the two tags (green) targetting both mitochondria and the ER and bridged by monomeric RFP. (**b**) Analysis of the expression of the linker targeted to the posterior compartment of the larval wing disc. Genotypes: hh>GFP, w;;hhGal4, UAS-GFP/+; hh>linker, w; UAS-linker/+; hhGal4, UAS-GFP/+.

### Linker expression increases MERCS and decreases mitochondrial length

Next, to determine if linker expression increases mitochondria-ER tethering, we quantified mitochondria-ER contacts in the neuronal cells of adult flies, as described previously ^3^. Ultrastructural analysis by transmission electron microscopy (TEM) revealed a significant increase in mitochondria-ER contacts in flies expressing the linker compared to controls (Fig. 2a and 2b). The points of contact between the ER and mitochondria correlate with the localisation of dynamin-related protein 1 (DRP1), a protein involved in mitochondrial fission in vertebrate cells. Such contact sites have been proposed to be involved in mitochondrial division (reviewed in ^1^). Confocal analysis of mitochondria labelled with a fluorescent tag (mito-GFP) in the mechanosensory neurons of the ventral nerve cord showed that the expression of the linker resulted in decreased mitochondrial length (Fig. 2c and 2d). Mitochondrial fragmentation in *Drosophila* has been proposed to drive the removal of defective mitochondria by mitophagy ^8^. To determine whether linker expression affects mitochondrial removal through fragmentation, we next assessed mitochondrial density by measuring the levels of the mitochondrial matrix enzyme citrate synthase, an indirect measure of mitochondrial density in flies ^9^. Citrate synthase levels in flies expressing the linker were not altered compared to those in controls (Fig. 2e), indicating that the mitochondrial fragmentation observed upon linker expression was not accompanied by a loss of mitochondrial mass. Mitochondria generate the majority of ATP as a source of cellular energy. We measured total levels of ATP in adult flies expressing the linker but did not find significant changes in ATP levels compared to those in controls (Fig. 2f). We also assessed the functional status of mitochondria by measuring the activity of respiratory complexes in flies expressing the linker and found no significant changes (Fig. 2g and 2h). Together, these results show that forcing contacts between mitochondria and the ER using an artificial linker induces mitochondrial fragmentation without altering global mitochondrial density or activity.

**Figure 2.**
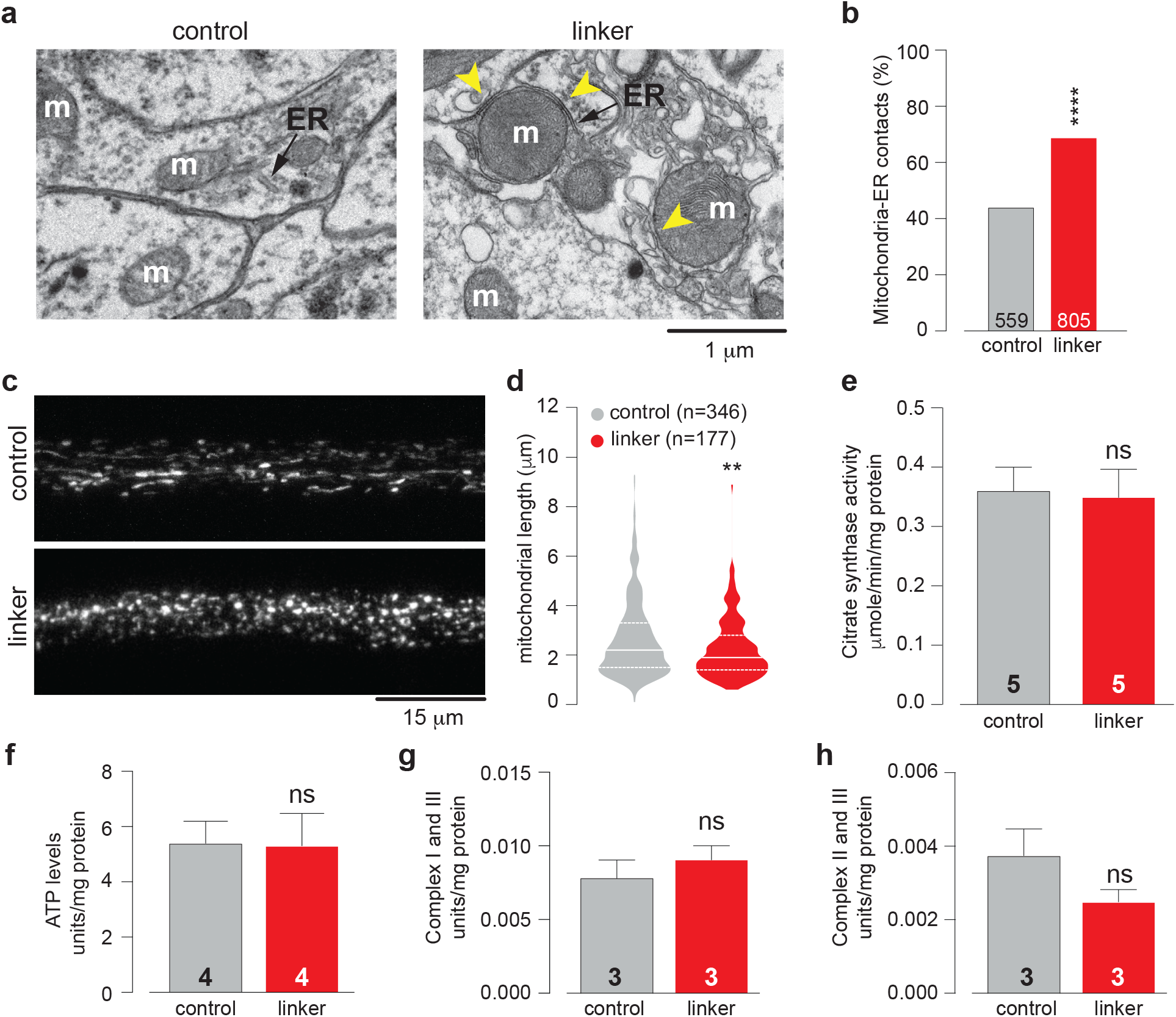
Forcing contacts between mitochondria and the ER induces mitochondrial fragmentation. (**a** and **b**) The quantification of mitochondria–ER contacts in adult fly brains (asterisks, chi-square two-tailed, 95% confidence intervals) (**b**) and representative electron microscopy images (**a**). The yellow arrows show mitochondria in contact with the ER (arrows). ER, endoplasmic reticulum; m, mitochondria. (**c** and **d**) The expression of the linker results in mitochondrial fragmentation. Confocal analysis of mitoGFP in the larval mechanosensory axons. Representative confocal images (**c**) and the quantification of mitochondrial length (**d**). The quantification of mitochondrial length is shown as a combined violin and box plot (p value, two-tailed unpaired *t*-test, compared to control). The expression of the linker does not affect overall mitochondrial mass. Mitochondrial mass assessed by measuring the activity of the mitochondrial matrix enzyme citrate synthase in adults (mean ± SD, ns, p>0.05, two-tailed unpaired *t*-test, compared to control). (**f** - **h**) The expression of the linker does not alter ATP levels (**f**) or mitochondrial function (**g**and **h**). Mitochondrial function was assayed by measuring the activity of NADH: cytochrome c reductase (**g**) and succinate: cytochrome c reductase (h) (mean ± SD, ns, p>0.05, two-tailed unpaired *t*-test, compared to control). Genotypes (**a** and **b**): control: w; elavGal4/+; +; linker: w; elavGal4/UAS-linker; +. Genotypes (**c**and **d**): control: w; elavGal4/+; UAS-mitoGFP/+; linker: w; elavGal4/UAS-linker; UAS-mitoGFP/+. Genotypes (**e**-**h**): control: w; +; daGal4/+, linker: w; UAS-linker/+; daGal4/+.

### Enhancing MERCS causes increased locomotor activity

We have previously shown that increased mitochondrial fission in flies with decreased expression of *Drosophila* mitofusin (*dMfn*) causes locomotor defects in adults^10^. To determine whether mitochondrial fission caused by linker expression affects motor activity, we measured locomotion in both larvae and adults by expressing the synthetic linker ubiquitously. First, we analysed locomotor activity in larvae using a motility tracking assay. We found a significant increase in the crawling speed (Fig. 3a) and total displacement (Fig. 3b) of larvae expressing the linker. Next, analysis of locomotor activity in adult flies for a period of up to 20 days showed a generalised increase in the activity of flies expressing the linker (Fig. 3c and 3d).

**Figure 3.**
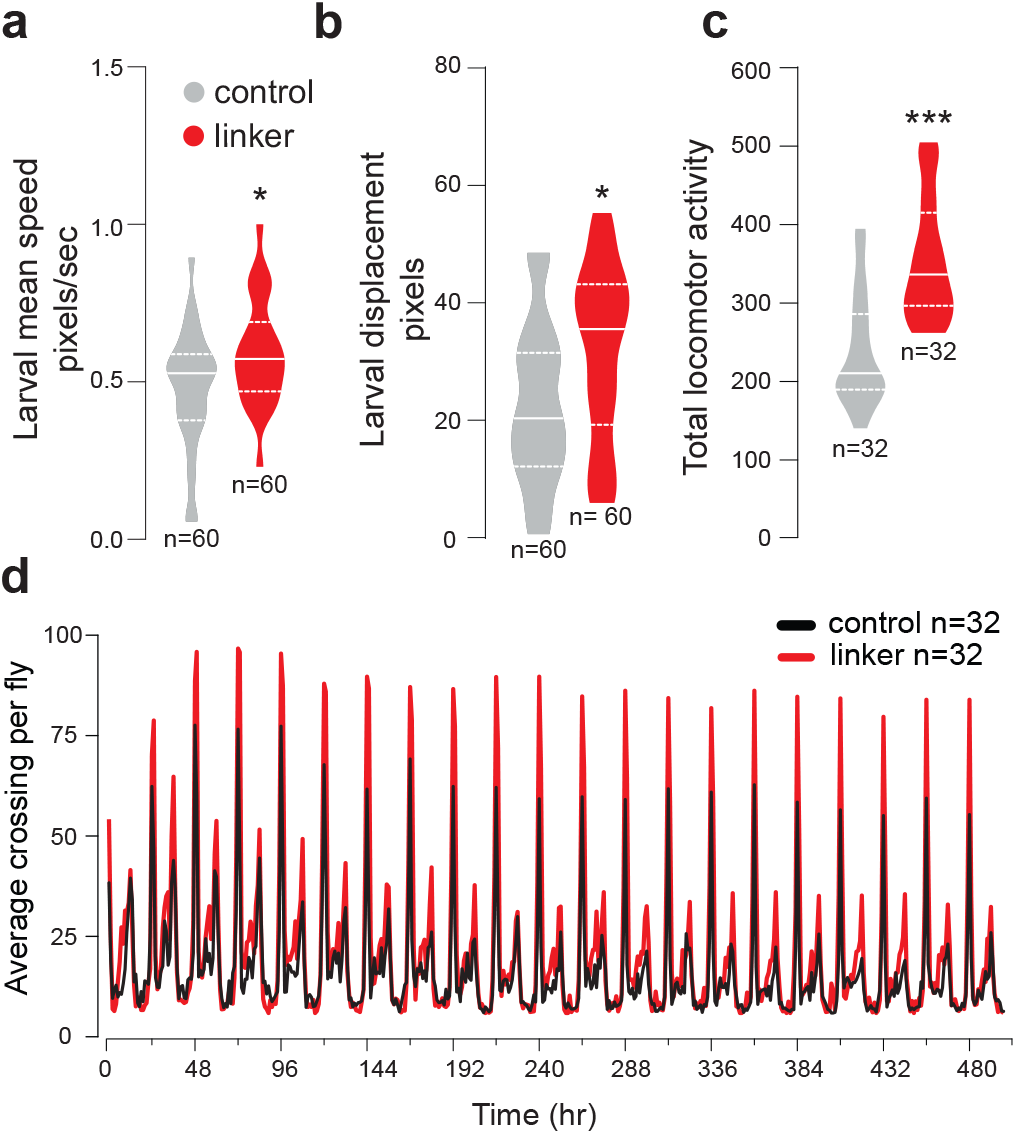
Enhancing MERCS increases locomotor activity. (**a** and **b**) Analysis of larval crawling in control and linker-expressing animals. Larval mean speed and displacement (**b**) were measured (mean ± SD; asterisk, two-tailed unpaired *t*-test). (**c**) Total activity in adult flies over a period of 20 days was measured using the Trikinetics system. (mean ± SD; asterisks, two-tailed unpaired *t*-test compared to control). (**d**) Actogram showing that the linker expressing flies have a higher average activity per hour than control over a period of 480 hours. Genotypes: control: w; +; daGal4/+; linker: w; UAS-linker/+; daGal4.

### Linker expression increased calcium levels in the mitochondria at ER contact points

MERCS play a key regulatory role in several cellular functions, including the transfer of Ca^2+^ and lipids between the ER and the mitochondria (reviewed in ^1^). Additionally, in hepatocytes, Ca^2+^ transport from the ER to mitochondria at contact sites elevates mitochondrial Ca^2+^ and increases mitochondrial ROS^2^. To measure Ca^2+^ levels in the mitochondria of flies expressing the linker, we used mitycam, a mitochondria-targeted fluorescent calcium reporter for *Drosophila* ^11^. As our artificial linker is tagged with RFP, we compared the mitycam signal in areas of RFP fluorescence (which correspond to mitochondria-ER contacts) to that in areas in which no RFP fluorescence was detected (which correspond to mitochondria distant from the ER). This revealed that mitochondria close to the ER contain higher levels of Ca^2+^ when compared to those in mitochondria distant from the ER (Fig. 4a and 4b).

**Figure 4.**
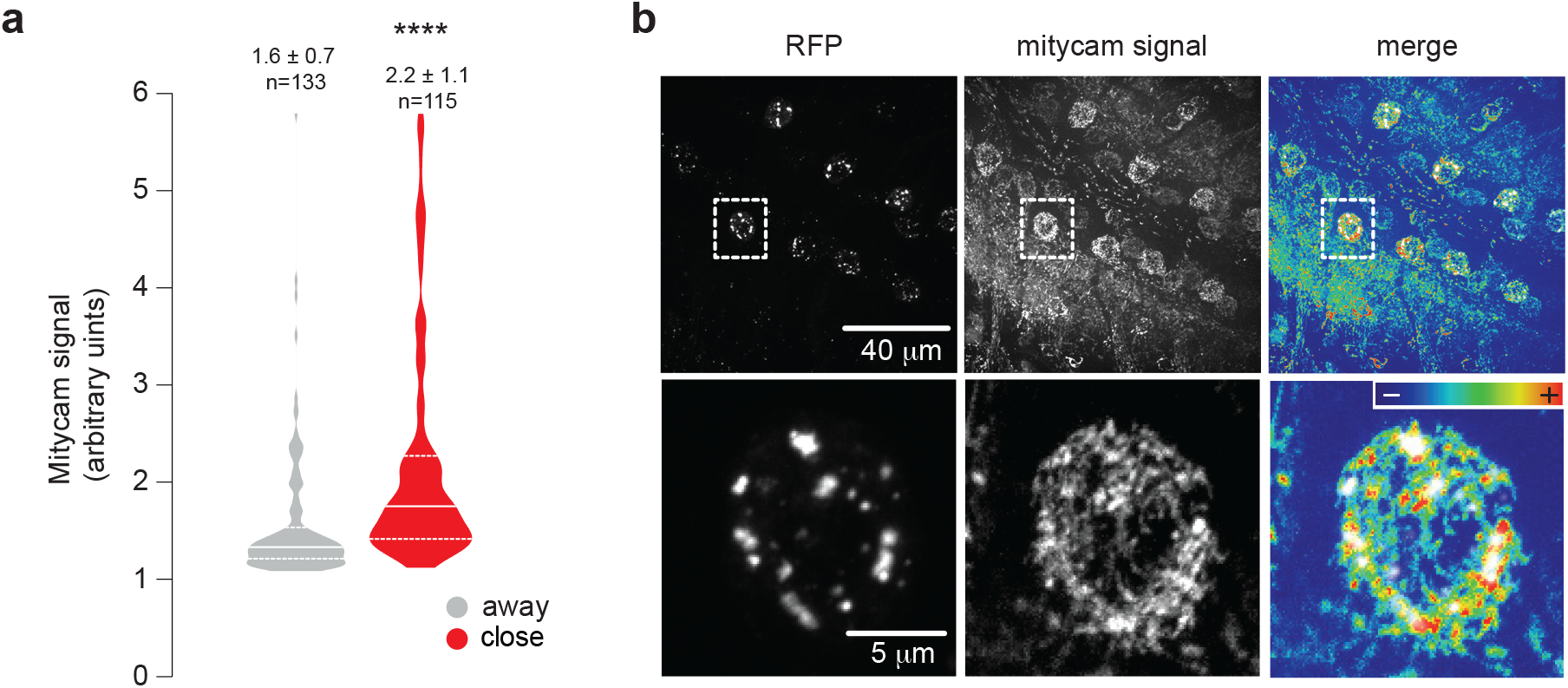
Increased calcium levels in mitochondria at MERCS. (**a** and **b**) Analysis of mitochondrial calcium uptake in neurons at the ventral nerve cord in larvae expressing the linker. The signal of the mitycam reporter (mitycam signal) was analysed close to and away from the linker signal (RFP). (**a**) The quantification of the mitycam fluorescence (mean ± SD; asterisks, two-tailed unpaired *t*-test compared to control, n = region of interest (ROI) measured). (**b**) Representative confocal images of the mitycam signal and the linker in larval brain cells. Genotypes (**a** and **b**): w; elavGal4, UAS-mitycam/UAS-linker; +.

### Artificially increasing MERCS increases ROS levels and enhances longevity

We next tested whether the linker expression affects the levels of mitochondrial ROS in adult flies. We performed this analysis using CellRox, a fluorescent ROS indicator. Our analysis showed an age-dependent increase in the levels of ROS (Fig. 5a and 5b) in adult flies expressing the linker. It has been shown that increased ROS production in flies extends lifespan ^12^. We therefore asked whether the effects on ROS levels induced by linker expression can affect the lifespan of adult flies. Our analysis showed that enhancing MERCS through linker expression extends lifespan (Fig 5c).

**Figure 5.**
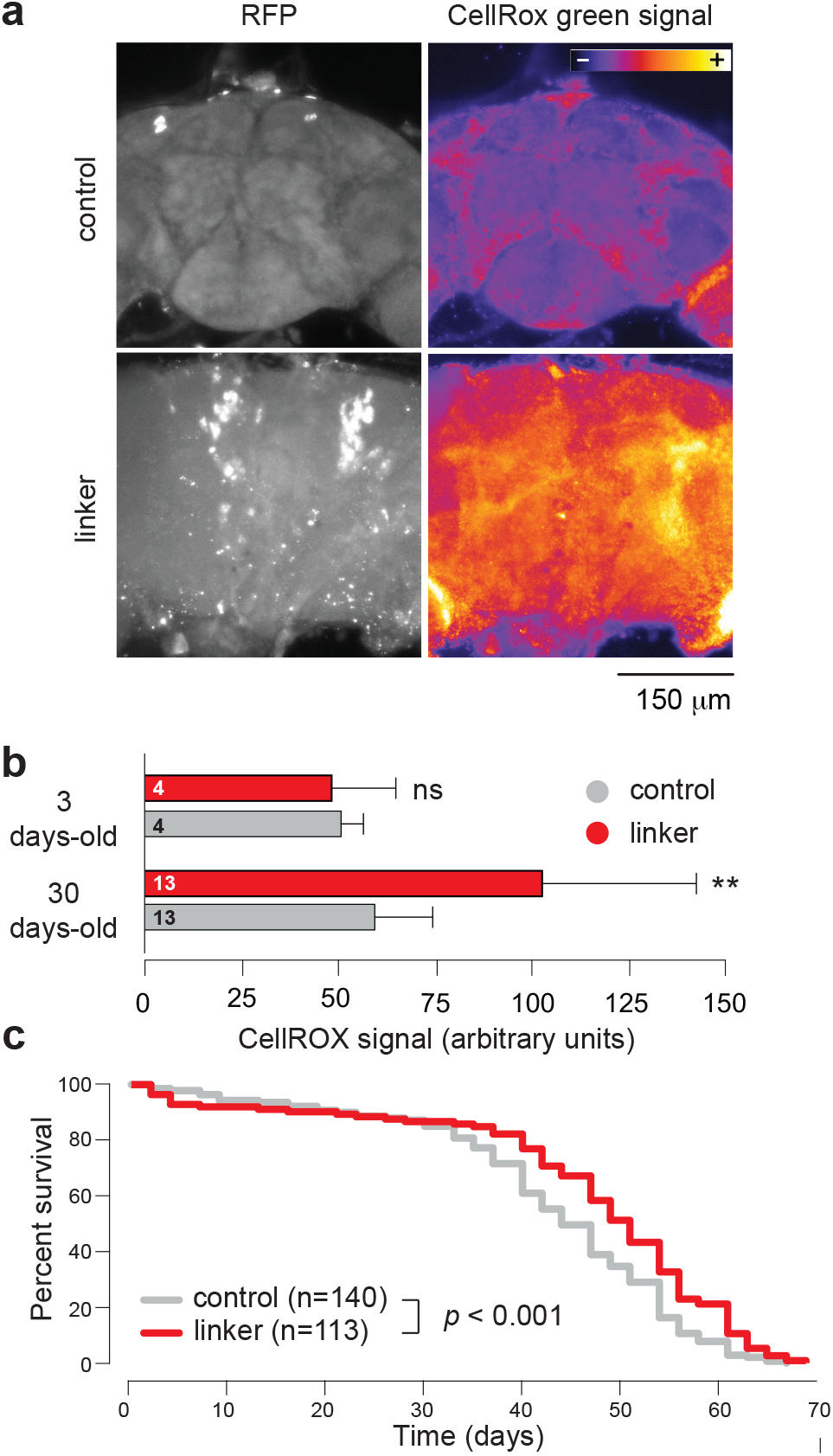
Enhancing MERCS increases ROS levels in aged flies and increases lifespan. (**a** and **b**) Analysis of mitochondrial ROS levels in young (3-day-old) and aged (30-day-old) controls and linker-expressing flies. Representative confocal images of 30-day-old brains stained with CellRox green are shown (**a**). The quantification of CellRox green signal (mean ± SD, asterisks, two-tailed unpaired *t*-test, compared to control) (**b**). (**c**) Lifespan analysis was performed over a period of 70 days (asterisks, log-rank Mantel-Cox test). Genotypes (**a**-**c**): control: w; +; daGal4/+; linker: w; UAS-linker/+; daGal4.

### Increasing contacts alleviates alterations in locomotion and lifespan in flies expressing *Aβ*_*42*_ *arc*

Signalling at mitochondria-ER contact points has been linked to several neurodegenerative disorders including Alzheimer’s disease (AD) ^13^. In AD, an abnormal toxic protein in the brain, amyloid-β (Aβ), accumulates and causes neuronal cell death. *Drosophila* can be used to model AD through the expression of arctic mutant (Glu22Gly) Aβ peptides (*Aβ*_*42*_ arc) in fly neurons. This causes progressive locomotor defects and the premature death of the flies ^14^. We therefore tested whether forcing mitochondria-ER contacts by linker expression affects the toxic consequences of the expression of *Aβ*_*42*_ arc in flies. We observed that linker expression suppressed climbing defects in flies expressing *Aβ*_*42*_ arc (Fig. 6a). Lifespan analysis confirmed that the decreased lifespan of flies expressing *Aβ*_*42*_ arc was suppressed by linker expression (Fig. 6b). We conclude that forcing mitochondria-ER contacts by the expression of the artificial linker confers protection in this *Drosophila* model of AD.

**Figure 6.**
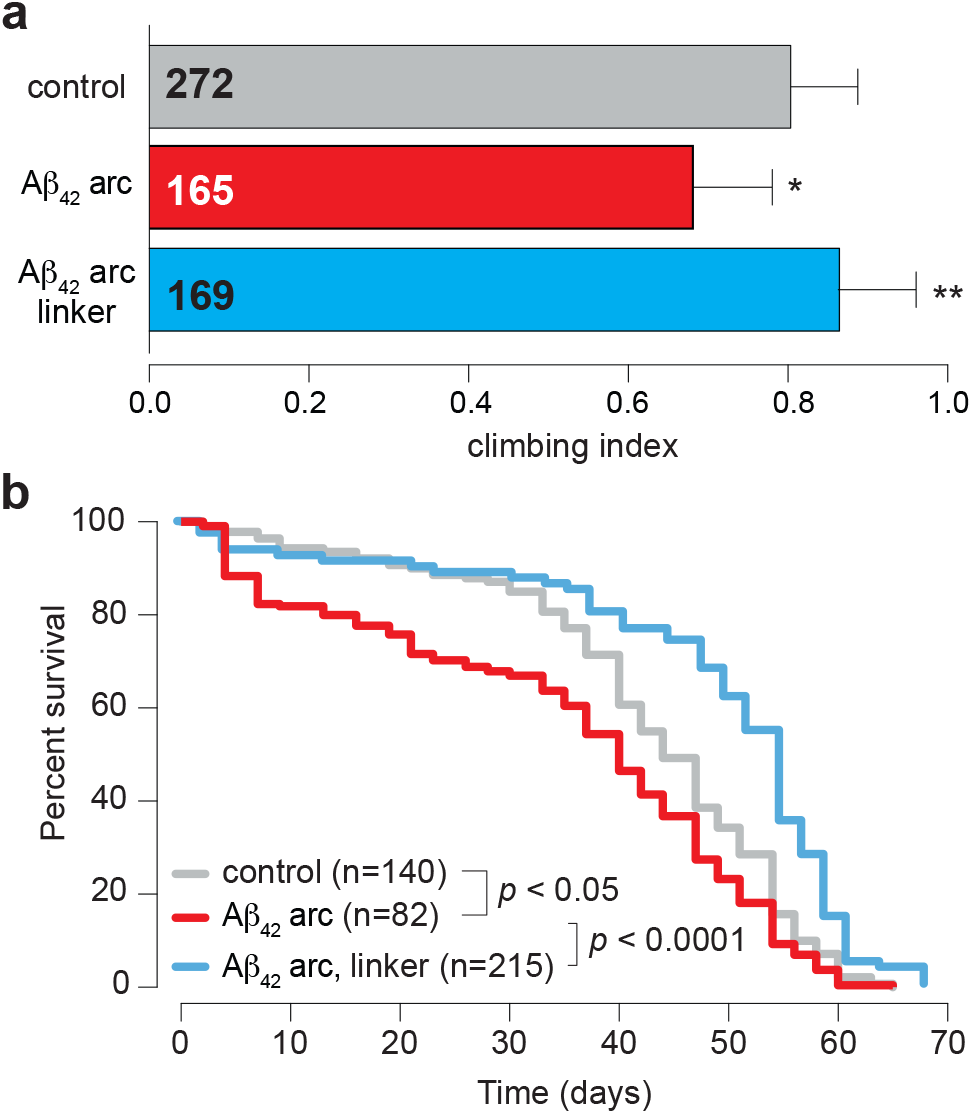
Linker expression rescues locomotor defects and increases lifespan in flies expressing toxic amyloid beta. (**a**) Analysis of locomotion in controls, Aβ42 arc flies and Aβ42 arc flies expressing the linker. Flies were tested using a standard climbing assay (mean ± SD, asterisks, one-way ANOVA). (**b**) Lifespan analysis was performed over a period of 70 days in controls, Aβ42 arc flies and Aβ42 arc flies expressing the linker (asterisks, log-rank Mantel-Cox test). Genotypes (**a** and **b**): control: w; +; daGal4/+; Aβ42 arc: w; +; daGal4/UAS-Aβ42 arc and Aβ42 arc, linker: w; UAS-linker/+; daGal4/UAS-Aβ42 arc.

## DISCUSSION

In this study, we show that linker expression leads to a decrease in mitochondrial length (Fig. 2c and 2d). The ER controls mitochondrial biogenesis, a process that involves mitochondrial division. The division of mitochondria through fission occurs at MERCS, in which the division machinery is recruited and mitochondrial constriction occurs (reviewed in ^1^). It is therefore possible that the decreased mitochondrial length observed in flies expressing the linker is a consequence of increased levels of mitochondrial fission resulting from a higher number of MERCS.

We detected increased levels of Ca^2+^ in mitochondria in close proximity to the linker (Fig. 4a and 4b). Increases in Ca^2+^ levels in the mitochondrial matrix activate enzymes involved in the citric acid cycle, including pyruvate, isocitrate and α-ketoglutarate dehydrogenases ^15, 16^ and stimulate oxidative phosphorylation, leading to an increased production of ATP. On the other hand, sustained levels of high Ca^2+^ in mitochondria are detrimental to their function (reviewed in ^17^). We did not observe any global alterations in ATP levels (Fig. 2f) or in mitochondrial function (Fig. 2g and 2h) upon linker expression. However, given that we measured ATP and mitochondrial function globally and not in the subset of mitochondria in close proximity to the ER, it is possible that local effects were diluted when the biochemical assays used in this study were performed.

Forcing mitochondria-ER contacts by linker expression in mice has been shown to lead to an increase in markers of ER stress and higher ROS levels. However, this has only been observed in animals subjected to overnutrition by a high fat diet ^2^. Here we detected a time-dependent increase in ROS levels in flies expressing an artificial linker without a specific dietary intervention. Historically, mitochondrial ROS have been associated with negative outcomes such as premature ageing ^18^ and diseases such as Parkinson’s disease ^19^. However, it has been shown that, in flies, increasing ROS production specifically through respiratory complex I reverse electron transport (RET) extends *Drosophila* lifespan ^12^. Additionally, this RET-mediated effect of ROS is capable of enhancing the lifespan of flies with decreased expression of *pink1*, a gene linked to Parkinson’s disease ^12^. Here, we found that linker expression was associated with higher ROS, enhanced longevity and the suppression of phenotypes associated with the expression of toxic Aβ. As toxic Aβ expression is linked to mitochondrial dysfunction (reviewed in ^20^), it would be important to determine if RET is the mechanism through which our artificial linker rescues the phenotype induced by toxic Aβ expression in flies. We propose that this could be investigated in future studies by epistasis analysis, for example, by expressing the alternative oxidase (AOX) of *Ciona intestinalis*, an enzyme that suppresses RET in flies ^12^. If ROS-mediated RET is the mechanism through which linker expression rescues mitochondrial dysfunction in flies expressing toxic Aβ, AOX expression should block this rescue.

In summary, by using an artificial linker to increase contacts between mitochondria and the ER, we uncovered how increased contacts between these two organelles can remodel signalling pathways with consequences for lifespan and disease outcomes.

## Abbreviations

ER: endoplasmic reticulum
MERCS: mitochondria-ER contact sites
Ca^+2^: calcium
AD: Alzheimer’s disease
Aβ: amyloid beta
MD: mitochondrial density
ATP: adenosine triphosphate
RFP: red fluorescent protein
OXPHOS: oxidative phosphorylation system
ROS: reactive oxygen species

## COMPETING INTERESTS

The authors declare no conflicts of interest.

## AUTHOR CONTRIBUTIONS

J.G.M. and S.H.Y.L performed the experiments and analysed the data. J.G.M., S.H.Y.L and L.M.M. wrote the paper.

## FUNDING

This work is funded by the UK Medical Research Council, intramural project RG94521.

## METHODS

### Genetics and *Drosophila* strains

Fly stocks and crosses were maintained on standard cornmeal agar media at 25°C. The linker (a gift from Dr. G. Csordás, Thomas Jefferson University, Philadelphia, USA), which encoded a monomeric red fluorescent protein (mRFP) preceded by the outer mitochondrial membrane targeting sequence mAKAP1 and followed by the ER targeting sequence yUBC6. This construct was codon optimised by site directed mutagenesis and cloned into the pUASTattB vector for PhiC31-mediated site-directed transgenesis. Transgenic flies were generated at the Cambridge fly facility, Department of Genetics, University of Cambridge, Cambridge, UK. *da*GAL4, *elav*-GAL4 and UASmitoGFP were obtained from the Bloomington Stock Centre, UAS-*Aβ*_*42*_ *arc* was a kind gift from D.C. Crowther, Department of Genetics, University of Cambridge, Cambridge, UK, and UASmitycam was a gift from Professor Julian Dow, Institute of Molecular Cell & System Biology, University of Glasgow, Glasgow, UK. All experiments on adult flies were performed with males.

### Citrate synthase assay

Citrate synthase activity was measured using the Citrate Synthase Assay kit (Sigma, CS070) as described previously ^10^. The units of citrate synthase were normalised to the protein concentration (mg/ml).

### OXPHOS activities

The activities of NADH: cytochrome c reductase (complex I + III) and succinate: cytochrome c reductase (complex II + III) were determined in fly extracts by spectrophotometry as previously described ^21, 22^ and normalized to the protein concentration (mg/ml).

### ATP levels

Five male flies (3- to 5-day-old) were homogenised in 100 μL of 6 M guanidine-HCl in extraction buffer (100 mM Tris and 4 mM EDTA, pH 7.8) to inhibit ATPases. Homogenised samples were subjected to rapid freezing in liquid nitrogen followed by boiling for 5 minutes. The samples were then cleared by centrifugation and the supernatant was diluted (1/50) with extraction buffer and mixed with the luminescent solution (CellTiter-Glo Luminescent Cell Viability Assay, Promega). The luminescence was measured on an Infinite M200PRO plate reader (TECAN, Switzerland). The relative adenosine triphosphate (ATP) levels were calculated by dividing the luminescence by the total protein concentration, which was determined by the Bradford method.

### Electron microscopy

For transmission electron microscopy, adult fly brains were fixed overnight in 0.1 M sodium cacodylate buffer, pH 7.4, containing 2% paraformaldehyde, 2.5% glutaraldehyde and 0.1% Tween-20. Then, the samples were post-fixed for 1 h at room temperature in a solution containing 1% osmium tetroxide and 1% potassium ferrocyanide. After fixation, the samples were stained with 5% aqueous uranyl acetate overnight at room temperature; then, they were dehydrated via a series of ethanol washes and embedded in TAAB epoxy resin (TAAB Laboratories Equipment Ltd., Aldermaston, UK). Semi-thin sections were stained with toluidine blue, and areas of the sections were selected for ultramicrotomy. Ultrathin sections were stained with lead citrate and imaged using a MegaView 3 digital camera and iTEM software (Olympus Soft Imaging Solutions GmbH, Mu nster, Germany) with a Jeol 100-CXII electron microscope (Jeol UK Ltd., Welwyn Garden City, UK).

### Larval crawling assay

For crawling analysis, 10 larvae were placed on a 92-mm square petri dish with 2% agar and monitored for 10 minutes under constant lighting and temperature. Video files were converted to images (1 frame per second) and analysed using ImageJ software. The images were imported as stacks and background-subtracted (average of all pictures). The TrackMate plugin ^23^ was used to analyse the movement of the larvae using displacement, speed and location parameters. Tracks lasting more than a minute were selected and used for data analysis.

### Lifespan analysis

Groups of 15 newly eclosed males of each genotype were placed into separate vials with food and maintained at 25°C. The flies were transferred to vials containing fresh food every 2-3 days, and the number of dead flies was recorded.

### Locomotor assay

Locomotor activity was recorded using the *Drosophila* Activity Monitoring System (TriKinetics, Waltham, MA, USA). In each assay, 32 1-day-old male flies of each genotype were grown under a light/dark 12 h:12 h cycle at 25°C. The number of average midline crossings per day were plotted for each genotype and analysed using GraphPad Prism. All experiments were performed in triplicate.

### Climbing performance

Climbing assays were performed using a counter-current apparatus equipped with 6 chambers. A total of 15 to 20 3-day-old male flies were placed into the first chamber, tapped to the bottom, and then given 20 s to climb a distance of 10 cm. The flies that successfully climbed 10 cm or beyond within 20 s were then shifted to a new chamber, and both sets of flies were given another opportunity to climb the 10-cm distance. This procedure was repeated a total of five times. After five trials, the number of flies in each chamber was counted. A video demonstrating this technique can be found at https://youtu.be/vmR6s_WAXgc. The climbing index was measured using a weighted average approach with the following formula:

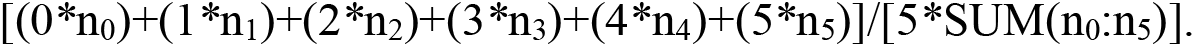

In this formula, *n*_*0*_ corresponds to the number of flies that failed the first trial, and *n*_*1*_ through *n*_*5*_ are the numbers of flies that successfully passed each successive trial. At least 100 flies were used for each genotype tested.

### Mitochondrial calcium imaging

Mitochondrial calcium was monitored by using a fluorescent mitochondria-targeted calcium reporter (mitycam). Mitycam is a modified pericam-based probe with a single excitation wavelength peak at ∼485 nm ^11^. For mitycam imaging, *Drosophila* larval brains were dissected in Schneider’s medium. Neurons expressing *UAS-mitycam* and *UAS-linker* driven by *elavGal4* were monitored using a Zeiss LSM510 confocal system. The reporter was excited with an argon 488-nm laser, and the emission signal was filtered through a 505-530-nm bandpass filter. Mitycam signals close to and distant from the linker signal (mRFP) were measured using ImageJ software.

### Measurement of mitochondrial ROS in Drosophila adult brain

Adult *Drosophila* brains were dissected in PBS and incubated with 5 μM CellROX Green Reagent, a fluorescent probe for measuring oxidative stress, (Molecular Probes, C10444) for 30 minutes. After incubation, the brains were washed with PBS for 10 minutes and imaged on a Zeiss LSM510 confocal microscope. A 100 μm-thick stack was acquired and used to measure the MitoSOX signal using ImageJ software.

### Statistical analyses

Statistical analyses were performed using GraphPad Prism 8 (www.graphpad.com). The data are presented as the mean values, and the error bars indicate ± SD. The number of biological replicates per experimental variable (n) is indicated in either the respective figure or figure legend. Significance is indicated as * for p < 0.05, ** for p < 0.01, *** for p < 0.001, and **** for p < 0.0001.

## ACKNOWLEDGEMENTS

We thank G. Csordás and J. Dow for reagents; the Bloomington Drosophila Stock Center for fly stocks; Fly Facility, Department of Genetics, University of Cambridge for generating the transgenic lines; T. Ashby and M. Patel for fly food preparation and T. Smith and M. Martin for their assistance with electron microscopy. We also thank G. Fedele for assistance with the TriKinetics data analysis.

